# Does local adaptation influence thermal responses in red coral populations across depth gradients? Transcriptomic insights for effective conservation

**DOI:** 10.1101/2025.05.12.653513

**Authors:** S. Ramirez-Calero, J. Garrabou, S. Suresh, M. Gut, M. Jou, X. Sarropoulou, P. López-Sendino, M. Zabala, JB. Ledoux

## Abstract

Marine heatwaves (MHWs) pose significant threats to marine biodiversity, including Mediterranean octocorals. Using a common garden experiment, we test whether differential transcriptomic responses to thermal stress between shallow and mesophotic populations of *Corallium rubrum* are shaped by their adaptation to the local environment, i.e. local adaptation. Six individuals from one shallow (15m) and one mesophotic (48m) population were exposed to control (18°C) and thermal stress (25°C) treatments, with samples collected at day 0 (T0), day 5 (T5), and day 10 (T10) for RNA sequencing (N=36). We revealed 1,957 differentially expressed genes (DEGs) in response to heat stress. Mild transcriptomic responses were observed in the shallow population (441 DEGs) characterized by heat shock proteins (HSPs) and developmental regulation. Conversely, a stronger and extensive response was observed in the mesophotic population with more than twice as many DEGs (1,081), predominantly associated with enhanced stress and wound healing mechanisms. Temporal transcriptional shifts were larger between T5 to T10 in the mesophotic population (1,497 vs 241), while more stable in the shallow (265 vs 271). Additionally, 172 DEGs, including HSPs, apoptosis and collagen, were found in the shallow population under control conditions, indicating transcriptional frontloading. The contrasting thermal stress responses between populations suggest distinct adaptative strategies potentially driven by local adaptation. These insights challenge the deep refugia hypothesis that considered mesophotic populations as potential sources for recolonization and active restoration of shallow populations threatened by MHWs. Our results support the need to integrate population-specific adaptive responses into conservation and restoration strategies for *C. rubrum*.

## INTRODUCTION

Corals dominating tropical and temperate reefs have experienced global declines owing to intense bleaching and mass mortality events related to anthropogenic climate change, particularly marine heatwaves (MHWs) (Hughes et al. 2010, 2017; Garrabou et al. 2022). Over the past decades, field surveys have revealed significant variability in the response of individuals and populations within the same species affected by these extreme events, while broad experimental efforts have facilitated the understanding of some of the factors driving these differences in thermal sensitivity (Grottoli et al. 2021). Building on experimental approaches such as common garden and reciprocal transplants, transcriptomics was used to investigate the differential gene expression underlying contrasted thermotolerance among individuals, leading to the definition of molecular phenotype (Reusch, 2013). For instance, using *in situ* reciprocal transplants, Bay and Palumbi (2017) demonstrated that stable gene expression patterns (i.e. low number of differentially expressed genes - DEG) were linked to better survival and growth in the coral *Acropora hyacinthus* in highly variable thermal environments. Furthermore, Kenkel et al (2013) showed, using common garden experiments, that different gene expression profiles maximize individual fitness of inshore populations of *Porites astreoides* compared to offshore ones. These studies provided two main outputs. First, the identification of key molecular mechanisms in corals response to thermal stress (i.e. enzymes, tumor necrosis factors, Rab/Ras and heat shock proteins) allowing to define a generalized coral environmental stress response (ESR) (Type A and B; Dixon et al., 2020). Such ESR can be used to classify the severity and extent of thermal impacts between populations. The second main output of these studies was to demonstrate the complex interplay between ecological and evolutionary processes shaping variability in transcriptomic responses to thermal stress, highlighting the importance of local adaptation and phenotypic plasticity (Palumbi et al. 2014). Disentangling the relative contributions of adaptation and plasticity is critical to understand and forecast the responses of coral ecosystems in the context of climate change (Torda et al. 2017).

Local adaptation refers to a population having higher fitness in its native habitat compared to any foreign population (Kawecki and Ebert 2004). By setting the focus on transcriptomes and molecular phenotypes, local adaptation drives genotype-by-environment interactions resulting in advantageous gene expression patterns (Savolainen et al. 2013; Sork 2017). For example, Kenkel and Matz, 2016 found that locally adapted inshore populations of *Porites astreoides* exhibit similar molecular phenotypes to their original environment after transplantation which enhances their thermal tolerance in comparison with offshore populations. Specifically, inshore corals had a larger capacity to regulate gene expression involved in the ESR. On the other hand, phenotypic plasticity corresponds to the organism’s ability to adjust its molecular phenotypes in response to environmental variability (Ghalambor et al. 2015). For example, plasticity responses of calcium signaling and focal adhesion gene expression have been characterized in *Seriatopora hystrix* with mildly bleached phenotypes displaying higher gene expression plasticity to maintain coral-symbiont relationships (Wang et al. 2024). In addition to phenotypic plasticity and local adaptation, transcriptional frontloading (Barshis et al. 2013), which is the pre-regulation of protective groups of genes, has been shown to improve individual’s thermotolerance by reducing the requirement for inducible stress response (Collins et al. 2021). This has been evidenced in scleractinian corals inhabiting contrasting environments in which rapid and large-scale frontloaded expression of certain genes (e.g., heat shock proteins, tumour necrosis factors - TRAF, immune-related genes), drove enhanced acclimation and resilience to heat stress and bleaching (Barshis et al., 2013; Bay & Palumbi, 2014; Brener-Raffalli et al., 2022; Savary et al., 2021; Vidal-Dupiol et al., 2022). Although knowledge on the molecular signatures of plastic and adaptive processes has greatly improved in tropical hexacorals (i.e. NOAA - bleaching response; Bay & Palumbi, 2014; Logan et al., 2014; Teneva et al., 2012), this remains largely unexplored for other groups of corals, particularly temperate habitat-forming octocorals (but see Pratlong et al. 2015; Beauvieux et al., 2024).

The Mediterranean Sea is known as a hot-spot of climate change, exerting impacts on marine biodiversity through mass mortality events (MMEs) associated to MHWs (Cramer et al. 2018; Garrabou et al. 2021, 2022). Over the past two decades, recurrent climate-driven MMEs impacted thousands of kilometers of coastal habitats across the Mediterranean, affecting around 90 species, across 8 phyla and driving local populations toward collapse (Garrabou et al. 2021, 2022; Gómez-Gras et al. 2021; CIESM 2023; Estaque et al. 2023). Habitat-forming octocorals such as *Paramuricea clavata* and *Corallium rubrum* are among the most affected species. These species exhibit a wide bathymetric distribution range from shallow to mesophotic and deep waters (Weinberg 1976; Knittweis et al. 2016). As such, shallow populations experience greater thermal variability, especially during the summer season, compared to mesophotic populations which are buffered from extreme thermal fluctuations. Moreover, populations in these species are usually genetically structured at low spatial scale (Ledoux et al. 2018; Gazulla et al. 2021). This combination of ecological variations and fine-scale genetic structuring suggests strong, and potentially contrasting, interactions between populations and their local environments with the potential to drive local adaptation and phenotypic plasticity in thermotolerance. Differential responses to thermal stress have been experimentally confirmed among Mediterranean octocoral populations (Crisci et al. 2017; Gómez-Gras et al. 2021, 2022; Ramirez-Calero et al. 2024; Rovira et al. 2024). However, a general picture regarding the underlying processes, and how they are influenced by local environmental factors such as depth and thermal regimes is still lacking. For instance, Gómez-Gras et al. (2022) implemented a common garden experiment in controlled conditions to investigate the thermal stress responses among populations from the same depth but distant geographic locations (>1,500 km) of *Paramuricea clavata*. All populations demonstrated high sensitivity to thermal stress, questioning the influence from local thermal regime. In contrast, common garden experiments involving red coral populations from the same geographic location but different depths (shallow vs. mesophotic), reveal highly contrasted thermal sensitivity. This suggested potential local adaptation, with shallow individuals better coping with thermal stress in comparison to individuals from mesophotic habitats (Torrents et al. 2008; Ledoux et al. 2015). Also, patterns of gene expression during thermal stress showed significant differences depending on the depth of origin of the individuals and their thermal history (Haguenauer et al. 2013; Pratlong et al. 2015). Although a few genes have been identified in relation to differential responses to thermal stress, further research is needed to better characterize the transcriptomic pathways involved, such as the generalized coral environmental stress response (ESR), in order to more accurately assess the extent and severity of temperature-induced damage, including distinctions between Type-A and Type-B stress responses (Dixon et al., 2020). An ESR Type-A would correspond to upregulated processes of cell death, reactive oxygen (ROS) species, NF-kappa B signaling, immune responses and protein folding genes promoting heightened stress responses, whereas Type-B would correspond to a downregulation of all these processes suggesting a reduced stress response. Moreover, while local adaptation has been suggested (e.g. Haguenauer et al. 2013), the occurrence of transcriptional frontloading remains to be tested. These gaps of knowledge are particularly worrying owing to their conservation implications beyond their direct implication on the response to marine heatwaves (Fifer et al. 2021). Notably, if differential processes between shallow and mesophotic populations are confirmed, the deep refugia hypothesis, which suggests recolonization of heatwave-impacted shallow populations by deeper populations (Gugliotti et al., 2019), and active restoration efforts sourced by mesophotic populations, should be reconsidered.

As such, this study investigates the molecular phenotypic responses of *Corallium rubrum* to thermal stress, combining a common-garden experiment with transcriptomic approaches focusing on two populations from contrasting environments: shallow vs. mesophotic. We aim to uncover generalized ESR strategies (Dixon et al. 2020), elucidate molecular signatures involved in coping with thermal stress across days and to test for transcriptional frontloading as a potential adaptive process to thermal stress (Barshis et al. 2013; Collins et al. 2021). To reach these objectives, we: i) characterize the overall early transcriptomic responses of *C. rubrum* to thermal stress; ii) compare the differential gene expression patterns to thermal stress between the shallow and mesophotic populations independently; iii) explore the transcriptomic variations across three time points between the two populations; and iv) compare the basal gene expression in control conditions between the two populations to investigate the occurrence of transcriptional frontloading. These molecular insights should inform conservation and restoration strategies, offering new perspectives for managing this species amid escalating climate change impacts.

## MATERIALS AND METHODS

### Model species

The precious red coral, *Corallium rubrum* (Linnaeus 1758), dwells in subtidal rocky habitats in the Mediterranean Sea and in the Eastern Atlantic Ocean (Zibrowius et al. 1984). It has been historically harvested for commercial purposes due to the high economic value of its calcitic skeleton used to manufacture jewelry (Tsounis et al. 2007). This species is a gonochoric and aposymbiotic coral characterized by low population dynamics (mean growth rate 0.25 ± 0.15 mm year^−1^; Garrabou and Harmelin, 2002; Marschal et al., 2004), late sexual maturity (10 years of age; Torrents et al., 2005) and low dispersal abilities (Ledoux et al. 2010; Gazulla et al. 2021). The most abundant populations are found along a wide bathymetric range from 10 to 120 m, although some populations can be found at depths of up to 1,000 m (Costantini et al. 2010; Knittweis et al. 2016). Thus, *C. rubrum* inhabits highly contrasted thermal habitats, which, combined with significant genetic differentiation at a low spatial scale, can potentially promote local adaptation to thermal regimes (Torrents et al. 2008; Ledoux et al. 2015).

### Sampling and experimental setup

Six adult colonies were randomly collected in October 2021 by scuba diving from two populations at different depths with contrasting thermal regimes along the Catalonian coast (Spain; Figure 1a). The first population was sampled (N=3 colonies) at 15 m depth at Cap Castell (3°13’10.233” N, 42°4’56.276” E), hereafter “shallow” population, while the second population was sampled (N=3 colonies) at 48 m depth at Pota del Llop (3°13’33.787” N, 42°2’52.736” E), hereafter “mesophotic” population. Each colony, representing individual genotypes, was retrieved in bags containing 2L of water and immediately transported in coolers to the Aquarium Experimental Zone (ZAE) of the Institut de Ciències del Mar (ICM-CSIC, Barcelona, Spain). Upon arrival, each colony was fragmented into six experimental replicates to be equally acclimated for one week in an open water system with 50 μm sand-filtered running seawater at 17-18°C (Figure 1). No mortality signs and tissue necrosis were detected during this period and all samples showed active polyps. After acclimation, the six experimental replicates from each individual were divided into two treatments (3 replicates each): a control (18°C) and a stress (25°C), following a common garden experimental set-up for ten days (Figure 1b). For the stress treatment, the temperature was increased stepwise from 18°C to 25°C over three days and maintained at 25°C until the end of the experiment. Colonies were fed during the entire experiment. Further details on the experimental setup can be found in Figure S1.

**Figure 1.**
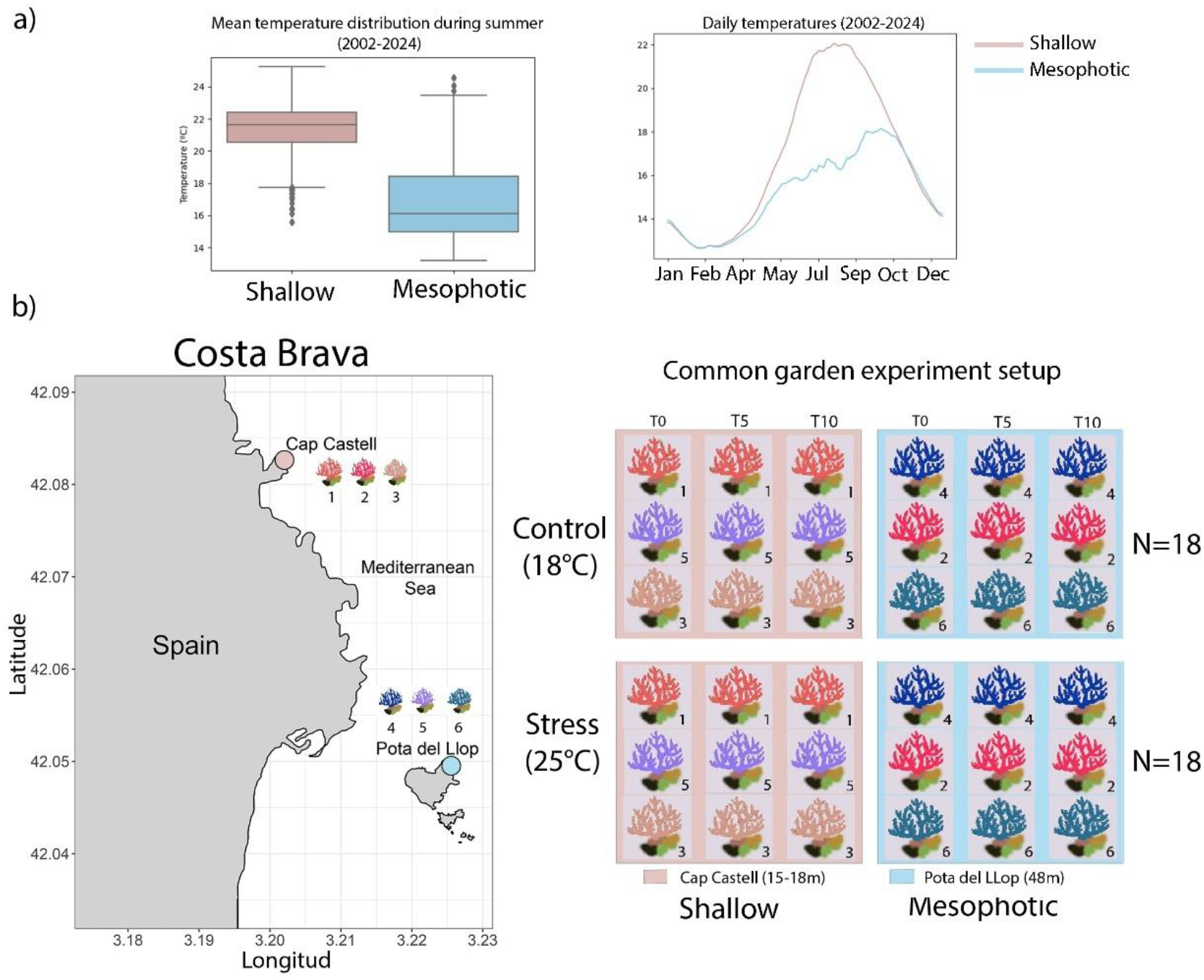
Annual thermal regimes characteristics, geographic location of sample sites, and experimental setup details for *Corallium rubrum*. a) Temperature records for shallow and mesophotic populations were obtained from the T-MEDNet database from 2002-2023 (Bensoussan et al. 2019). Plot lines are based on mean daily temperatures filtered by a running average, using 15 days as reference (360 records). b) Sample collection sites: Cap Castell (shallow population, 15-18m) and Pota del Llop (mesophotic population, 48m). c) Common-garden experimental design with colonies of *C. rubrum* exposed to control (18°C) and stress treatment (25°C). Each Individual (genotype; indicated by different colors and numbers) was randomly replicated across the experimental conditions and sampled at three time points: T0, T5, and T10

**Figure 2.**
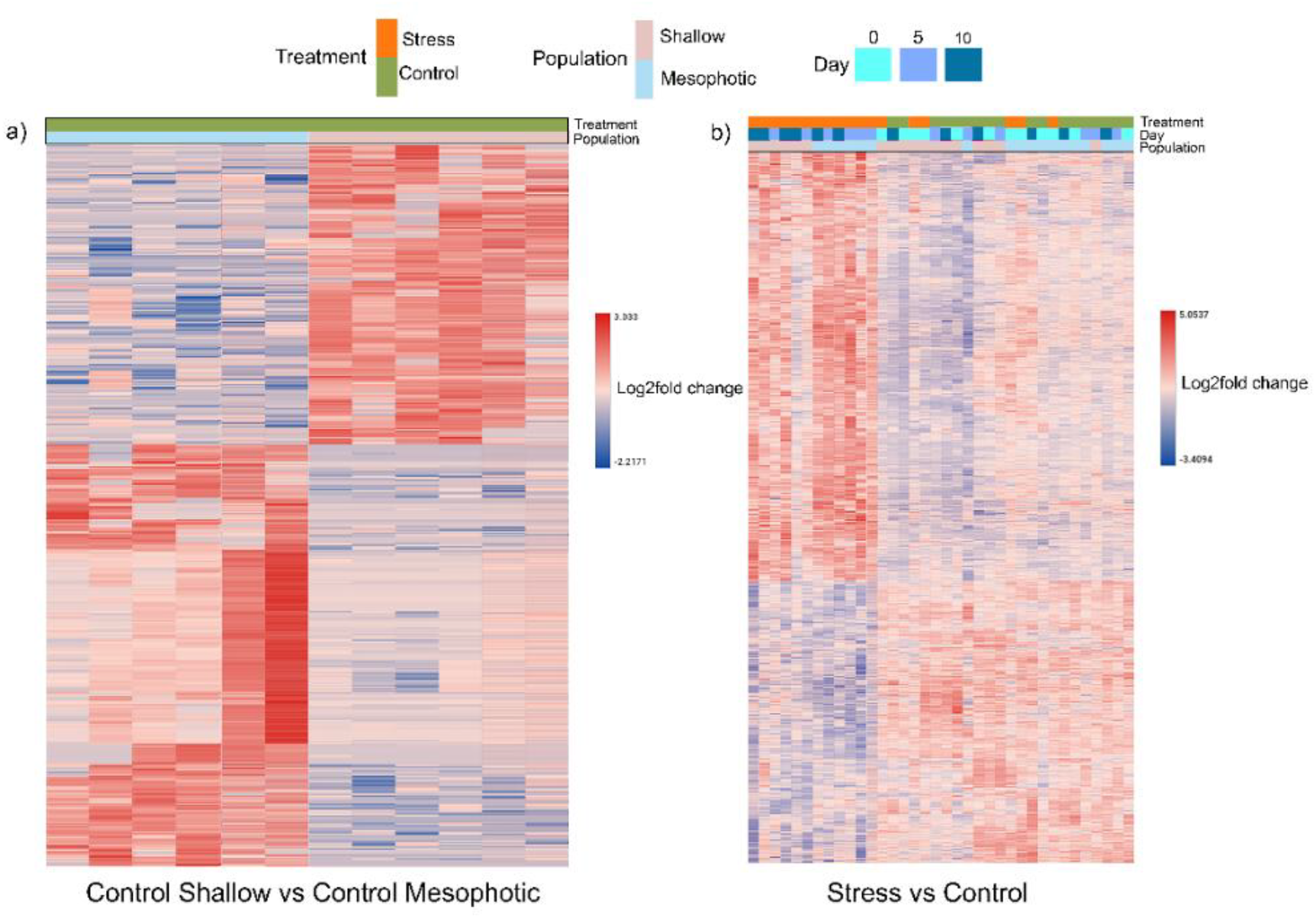
a) Differential gene expression patterns for baseline conditions (control samples + individuals at T0) between populations; b) Differential gene expression patterns for both populations during the experiment. The legend indicates the reference values of log2fold changes (blue – downregulated, red - upregulated) for each significant DEG found under each comparison. Figure references are represented in colors, populations being highlighted in pink (Shallow) and light blue (Mesophotic), conditions in green (Control) and orange (Stress), while days are in cyan (T0), cobalt (T5), and deep sky blue (T10). Further details can be found in Supplementary Tables S3a, b and S4a, b

### Thermal regime characterization

Temperature data were obtained from the T-MEDNet database (Bensoussan et al. 2019, Jou & Ramirez-Calero, 2024), a continuous monitoring effort of sea water temperature since 2002 across several regions in the Mediterranean Sea (Bensoussan et al. 2019). Several descriptors were used to characterize the thermal regimes of the two sampling locations. Mean daily temperatures of the hourly records were obtained after applying a running average filter of 15 days (360 records) in order to smoothen the data (Figure 1a, Table S1a). Standard deviation (SD) was calculated using the filtered data to assess its variability. Since the data presented some gaps in some records, the filtering was treated within blocks to disconnect distant dates. This filter helped reveal the differences in temperature variation between the two sites with a temporal perspective (Figure S2b). Accordingly, the shallow population (15m) was characterized by a mean maximum temperature of 24.19°C ± 1.85 SD, following the 2002-2023 records (Figure S2a, Table S1a). During the summer months (July, August, September - JAS; Hobday et al., 2016), the mean temperature corresponded to 22.54°C ± 1.66 SD, while the maximum temperature record reached 26.30°C (Table S1b). Considering marine heatwaves lasting more than five consecutive days during the summer months, the maximum temperature reached 24.44°C (Table S1c). Extreme heat days surpassing ≥ 23°C, 24°C, and 25°C during the summer season were more common in the shallow population (Table S1b). For the mesophotic population (40m), the mean maximum temperature corresponded to 22.83°C ± 1.82 SD (Figure S2a, Table S1a). During the summer months, the mean temperature corresponded to 17.79°C ± 2.22 SD, while the maximum temperature reached 25.09°C during this season (Table S1b). Marine heatwave events, with a duration of more than 5 consecutive days, reached a maximum of 23.56°C (Table S1c). Finally, only records of extreme heat days of ≥ 23°C, and one of ≥ 24°C were found in this population (Table S1b).

### RNA extractions

Fragments of *C. rubrum* were sampled threefold for RNA sequencing during the initial day of the experiment (T0), day 5^th^ (T5), and day 10^th^ (T10) in the two treatments (control and stress) and stored in RNAlater at -20°C (Figure 1b). RNA extractions were conducted using the RNAeasy Mini Kit protocol (Qiagen) with a modification of the lysis step that involved smashing the tissue with a sterile cane in TRIzol™ (Thermo-Scientific; see supplementary material). The resulting RNA was quantified using the Qubit® RNA BR Assay Kit (Thermo Fisher Scientific), while quality was assessed using the RNA Integrity Number (RIN), obtained on a Fragment Analyzer system, using the DNF-471 RNA kit (Agilent). Samples with a RIN > 6 were retained. mRNA-focused sequencing libraries were prepared using the KAPA Stranded mRNA-Seq Kit for Illumina Platforms (Roche), following manufacturer’s recommendations, starting with 500 ng of total RNA. Illumina-compatible adapters with unique dual indexes and unique molecular identifiers (Integrated DNA Technologies) were ligated to the RNA fragments. The ligation products were enriched via 15 cycles of PCR. The final libraries were validated using the DNA 7500 Assay on an Agilent Bioanalyzer 2100 system. Sequencing was performed on the NovaSeq 6000 platform (Illumina) in paired-end mode with a read length of 2 × 151 bp, generating 40–60 million reads per library. Sequencing followed the manufacturer’s protocol for dual indexing. Image analysis, base calling, and quality scoring were processed using Real-Time Analysis (RTA) software version 3.4.4, with FASTQ files generated as the final output.

### De novo *Transcriptome assembly*

Raw read quality was examined using FastQC v0.11.7 (Andrews 2010), then reads were trimmed and adapters were removed using Trimmomatic v0.38 (Bolger et al. 2014) with parameters as follows: ILLUMINACLIP:TruSeq3-PE.fa:2:30:10 LEADING:4 TRAILING:3 SLIDINGWINDOW:4:15 MINLEN:40. Potential fungal, bacterial, and viral contamination was removed with Kraken v 2.0.9-beta (Wood et al. 2019) using the standard database and a confidence score of 0.3. At the time of the study, no genome was available for the species, so we designed a *de novo* transcriptome assembly using reads from all six colonies with the trinity software v 2.13.2 (Haas et al. 2013), using the following parameters: *-- seqType* for fastqc data, a *--max_memory* 20G, *--full_cleanup, --SS_lib_type*, and *--left --right*. Assembly statistics were obtained using trinity script *TrinityStats*.*pl*.

To assess the quality of the assembly, we revised the read representation against our transcriptome using the aligner Bowtie2 v2.4.3 with default parameters (Langmead and Salzberg 2012). To reduce transcript redundancy and to obtain only the longest coding transcripts (>100bp), we used transdecoder v5.5.0 (Haas, BJ. https://github.com/TransDecoder/TransDecoder) to identify candidate ORF (Open Reading Frame) regions by keeping the option *– single_best_only*. From these results, we retrieved the output file to conduct a BLASTX analysis using the curated Swissprot/Uniprot database (Bateman et al. 2021). We addressed the completeness of our assembly using BUSCO 5.4.6 (Simão et al. 2015) to obtain the number of conserved orthologs content from our results represented in the dataset *metazoan_odb10*. Finally, we computed the Nx statistics to estimate the approximate length of our transcripts using the N50 statistics from the *trinity*.*stats*.*pl* script. This allowed us to determine the specific length of at least half of our transcripts, thus evaluating their contiguity and quality after the assembly process. Further transcriptome assembly metrics and details can be found at supplementary materials (Figure S3 and Table S2).

### Annotation

We annotated the transcripts of our transcriptome reference with BLAST+ (Camacho et al. 2009) using the curated Swissprot/Uniprot database and a custom database with curated genome annotations from coral species available in ENSEMBL: *Acropora millepora* (GCA_013753865.1), *Dendronephthya gigantea* (GCA_004324835.1), *Orbicella faveolate* (GCA_002042975.1), *Pocillopora damicornis* (GCA_003704095.1), *Stylophora pistillata* (GCA_002571385.1) and *Paramuricea clavata* (GCA_902702795.2). These were the closest species to *C. rubrum* with available genome annotations. A custom database was built using the function –*makeblastdb* with default parameters. Finally, we used Omicsbox v3.1.11 (BioBam Bioinformatics 2019) to functionally annotate the transcripts with Gene Ontology (GO terms) and associated pathways.

### Differential gene expression analyses

We first quantified the abundance of transcripts available in our reference using the trinity script *align_and_estimate_abundance*.*pl*, as follows: i) reference was built with *--est_method RSEM -- aln_method* bowtie2 *--trinity_mode --prep_reference* and ii) quantification was run by indicating only RSEM v1.3.3 as quantification method and Bowtie2 as mapping tool in the same way as for the reference. Final output files were used to build a gene expression matrix using the script *abundance_estimates_to_matrix*.*pl*. Transcripts with no expression were removed using the script *filter_low_expr_transcripts*.*pl*, retaining only the most highly expressed isoforms for each gene using the parameter *–highest_iso_only*. Before determining the differential gene expression of our samples, we performed a principal component analysis (PCA) using the R-package *pcaExplorer* v2.16.0 (Marini and Binder 2019). Further details can be found in supplementary information (Figure S4).

To evaluate differential gene expression given by the temperature stress, we used the DESeq2 package v1.30.1 (Love et al. 2014) in R, using a likelihood ratio test (LRT) with a model comparison approach to determine the effects of treatment (control vs. stress), population (shallow vs. deep), and time (days – T0, T5 and T10) on the gene expression patterns. To determine the best design formula for the final differential expression analysis, we measured significant differences by comparing a full model including treatment, population, and time against three reduced models including i) population and time (effect of temperature), ii) population and treatment (time effect), and iii) treatment and time (population effect). For a total of 3,192 DEGs due to population effect, 1,781 DEGs due to temperature effect, and 40 DEGs due to time effect, the factors that better explained the observed differences in gene expression corresponded to the reduced models including treatment and time (population effect) and population and time (treatment effect). Consequently, Wald test statistics setting a *p-adj* < 0.05 were implemented as follows: i) overall baseline gene expression between populations including only control treatment and samples from stress treatment at T0 (*design = ∼population*) as no effects of temperature were present at this time point; ii) overall effect of stress vs control for the entire data-set (i.e. including all samples) using the formula *design = ∼population + treatment + day*, iii) effect of stress independently for each population (i.e. gene expression detected only for one population) using the formula *design = ∼treatment + day*, and iv) effect of stress at each time point (T0, T5 and T10) for each population *design = ∼factor*. Further filters were set to avoid false positives such as absolute log2fold change (log2FC) greater than 0.3 and a baseMean greater than 10 as suggested in DESeq2 manual (Love et al., 2014).

Gene Ontology (GO) functional enrichments for significant differentially expressed (DE) genes were performed using Fisher’s exact test in Omicsbox v3.1.11 (BioBam Bioinformatics 2019), with a False Discovery Rate (FDR) threshold of 0.05, using the option “reduce to most specific” to avoid redundancy in the enriched GO terms identified. Heatmaps showing relative transcript expression for each test across samples were created using the function *pheatmap* v 1.0.12 in R (Kolve 2019) and Omicsbox. Additional figures were created with ggplot2 package v3.3.5 I (Wickham 2016).

### Transcriptional frontloading

We tested the transcriptional frontloading hypothesis (Barshis et al., 2013) between shallow and mesophotic populations. Our aim here was to test if the shallow population, which during summer is exposed to higher temperatures, may display a frontloading strategy conferring advantage to thermal stress over the mesophotic population, usually exposed to more stable thermal conditions (4.74 degrees colder; Table S1a, b). Accordingly, we assessed whether the significantly upregulated DEGs found in the mesophotic population under thermal stress were already upregulated in the shallow population in control conditions. Based on this, we categorically designate such upregulated genes as constitutively “frontloaded”. In addition, genes with a diminished log2fold change in response to temperature, meaning that gene expression levels under control do not erratically change their transcription during treatment, were considered as DEGs presenting a “reduced reaction” to temperature. Finally, DEGs labelled as “greater log2fold change” referred to genes having a larger proportional log2fold change difference when comparing baseline (control) expression to expression levels after exposure to thermal stress. Thus, a more pronounced response in gene expression is observed, reflecting the capacity of the shallow population for rapid amplification of stress-related functions. All definitions used here followed previous studies also assessing frontloading in corals (Barshis et al., 2013; Brener-Raffalli et al., 2022; Collins et al., 2021; Vidal-Dupiol et al., 2022).

## RESULTS

### Overall transcriptomic landscape for Corallium rubrum in control conditions

To understand the baseline differences between *C. rubrum* populations, we compared shallow vs mesophotic colonies under control conditions. By combining all control and stress replicates at day 0 (T0), a total of 189 DEGs were differentially expressed between the two populations (Table S3a). Such DEGs were involved in cellular structural processes and maintenance as well as metabolic processes such as lipids, fatty acids, and mitochondrial functions. No DEGs were related to stress or apoptosis. The major GO term categories confirmed that the majority of the DEGs belonged to cellular processes including ribosomes, endoplasmic reticulum, and cytoplasmic translation (Table S3b).

Additional differential gene expression analysis under control conditions, conducted independently for each population, revealed only three DEGs in the shallow population and 38 DEGs in the mesophotic population (Table S3c-S3d). Among the three DEGs in the shallow population, only one had an available annotation, corresponding to a member of the solute carrier family 49 (*SLC6A20*; Table S3b). Among the 23 DEGs in the mesophotic population with a valid annotation several transcriptional factors (elongation, splicing) and ribosomal proteins were found (*RpL32, RPS12, FAU, TCF15, Isy1, eef1b*, Table S3d).

### Differential effects of thermal stress for Corallium rubrum

To have an overall view of the thermal stress response in *C. rubrum*, we compared all replicates from the stress treatment against all control replicates, regardless of population. From this, a total of 1,957 DEGs were significant during treatment when considering the entire dataset, with 1,029 upregulated and 928 downregulated genes (Table S4a). From these DEGs, upregulated processes of apoptosis (*CRP3, ube2z, app*), immune responses (*Arl1, FEZF2, arf1*,) and hypoxia (*Chchd2, HIF1A, Egln1, hif1an*) were detected. In addition, stress signatures (*CIRBP, Afg1l, CIRBPB, Aco2*), heat shock proteins (*HSPA6, Dnajb9*) and developmental activity (*Cux1, GPSM2, Lrrc7*) were differentially regulated across the treatments (Table S4a). Major GO terms enriched among the DE genes were tumor necrosis factors, inflammatory responses, reactive oxygen species (ROS), hyperoxia and response to heat (Table S4b).

### Population specific gene expression patterns during thermal stress

To further characterize population specific responses to thermal stress, we first subset the data for each population, and then compared their respective stress treatment replicates against their control counterparts. Accordingly, the shallow population revealed fewer molecular alterations (441 DEGs; 240 upregulated and 201 downregulated) in comparison with the mesophotic (1,081 DEGs; 529 upregulated and 552 downregulated) during thermal stress, with a total of 199 DEGs shared between both populations (Figure 3a, Table S5a). Major GO term categories of such shared responses revealed that the majority of these shared DEGs belonged to developmental processes, oxygen homeostasis (hypoxia), metabolism and apoptosis (Table S5d).

**Figure 3.**
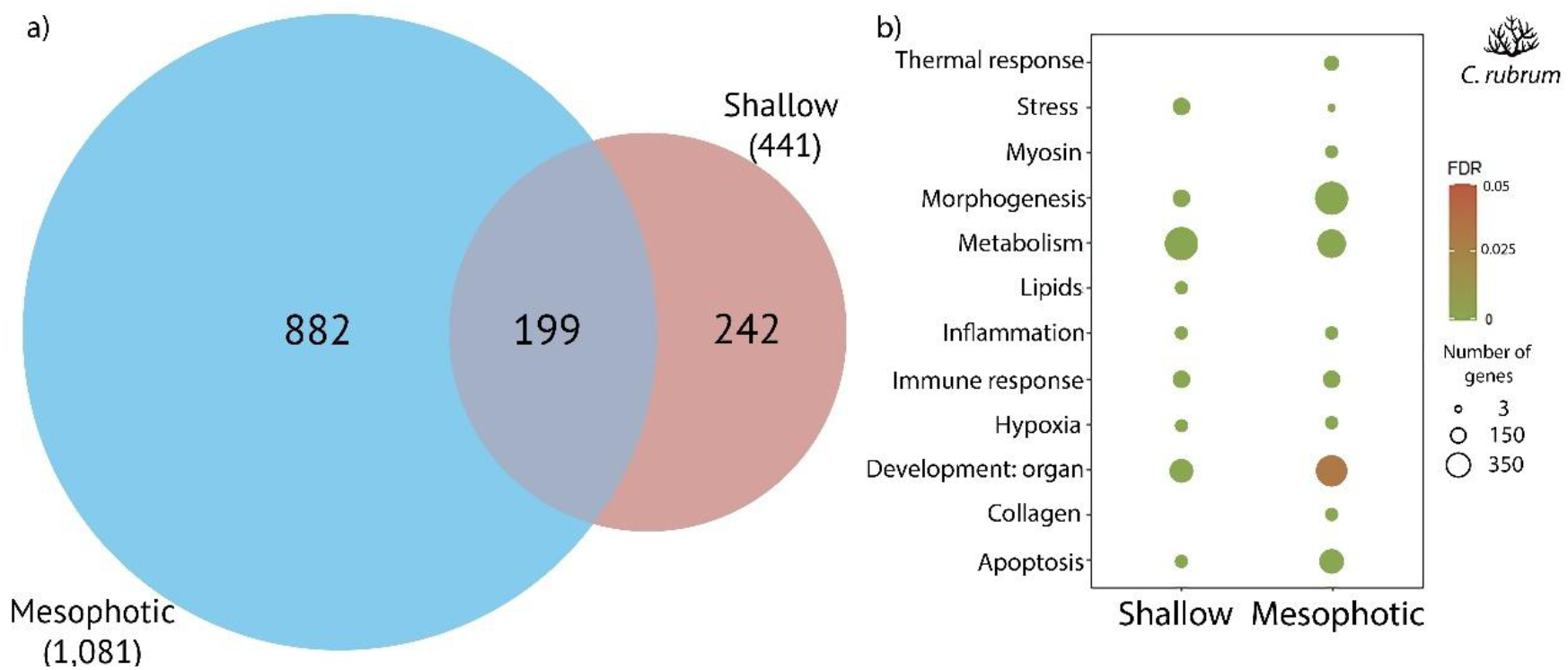
a) Number of differentially expressed genes between control and stress treatment for each population and the common overlap. b) Functional enrichment of Gene Ontology (GO) terms in the two populations of *C. rubrum* upon thermal stress. The size of the circles is proportional to the number of genes observed within each GO category, and the color of the circles is proportional to the significance (FDR value < 0.05)

The specific DEGs observed in the shallow population (242 DEGs; Figure 3a) suggested only one transcriptional process of apoptosis underlined by upregulated tumor necrosis factors (*Tctp, AIFM2*) and cell proliferation genes (*FGFR4*). DNA damage and stress signatures highlighted the expression of transcriptional repressors (*hes1-b, HNRNPK*), heat shock proteins (*HSP90B1*), inflammatory mediators (*Smpd1*), and autophagy enzymes (*Atg10*) (Table S5a). The significant expression of glycoproteins (*Clec4g*) and ubiquitin (*CYLD*) also revealed innate immune responses. Major GO term functions confirmed several processes of development altered by the up and down-regulation of specific muscle development genes (*Klhl40*), several enzymes (*PRKAA2, FDPS, HMGCR*), and hox genes (*EMS*) (Table S5a, b). Finally, significant DEGs were found under GO categories of hypoxia, inflammatory response, and metabolism (i.e. lipids, fatty acids oxidation) in this shallow population (Figure 3b; further details in Table S5a, b).

The mesophotic population exhibited larger unique transcriptomic alterations underlined by 882 DEGs (Figure 3b, Table S5a). For instance, a higher number of upregulated genes of tumor necrosis factors (*ZFAND6*), cell death-associated proteins (*htra1, Dapk3*), apoptosis repression and regulation (*tax1bp1b, STAT5A, MSX2*) highlighted the main differences in apoptosis-related processes. Also, genes involved in different processes of stress were transcribed, including actin stress fibers (*SORBS2, limch1*), oxidative stress genes (*chmp5, ACO2, PON2, FCN2*), proteotoxic stress (*RPS27A*), and endoplasmic reticulum stress – ER stress (*ATF6*) (Table S5a). Further DEGs were related to maintenance of developmental and morphogenesis processes (*PCDHY, ACT1, IFRD1)* (Figure 3b, Table S5a) and molecular alterations in immune and inflammatory responses were characterized by the transcription of specific enzymes (*senju, Nagk*). In particular, the transcription of heat shock proteins (*Hspa8*), cell death genes (*htra1*), aquaporins (*Aqp8*), endoplasmic reticulum stress genes (*DNAJB1, DNAJC3*) characterized the GO term of “response to temperature stimulus” (GO:0009266), only found in this population. Finally, further GO term categories revealed cellular structure processes related to collagen, myosin and fibrillin with up- and downregulation of DEGs related to morphology structure (*MYL9, MYLK*), muscle regulation (*sqh*), several collagen proteins (*COL2a1, COL12A1, COL6A2*,), and elasticity (*Fbn2, Fbn1*). These processes were also uniquely expressed in the mesophotic population (Figure 3b; Table S5a, c).

### Temporal dynamics of early thermal stress response during days 0, 5 and 10

To characterize the dynamics of early thermal stress responses in *C. rubrum*, we compared replicates from each time point against their respective controls (i.e. Shallow_Stress-T5 vs Shallow_Control-T5), independently for each population. Accordingly, for the shallow, no DEGs were significantly transcribed for replicates at T0. However, at T5, 265 unique DEGs were significant (142 upregulated and 123 downregulated; Figure 4a, Table S6a). These DEGs were related to apoptosis (*Litaf, COMP*), histone regulation (*SSRP1*), hypoxia inducible factor 1a (*HIF1A*), as well as endoplasmic reticulum (ER), genotoxic and oxidative stress (*BRSK2, cirbp-b, erm*). Developmental processes were significantly transcribed with major homeobox transcription factors (*<u>Uncx</u>*), myosin (*Mhc*), and collagen (*COLL7*) (Table S6a). Furthermore, at T10, the largest differential gene expression was revealed with 271 unique DEGs (123 upregulated, 148 downregulated; Figure 4a, Table S6a). Significant DEGs related to oxidative stress (*Bdh2*) as well as heat shock proteins (*ST13*) were upregulated. Finally, several genes related to transcription (*RPS25, RPS17, RPS23*), immune response (*Samhd1*), and microtubules (*GPSM2*) presented the largest changes in expression (|log2fold>17|) with respect to control conditions (Table S6a). Major enriched functions revealed diverse metabolic processes and responses to stimulus, while only one GO term was enriched for apoptosis, underlined by DNA damage genes (*MDM2*) and homeobox proteins (*EMX2, EMS*). No further GO terms related to stress or processes related to temperature were found in any of the days (Figure 4c, Table 6b, c).

**Figure 4.**
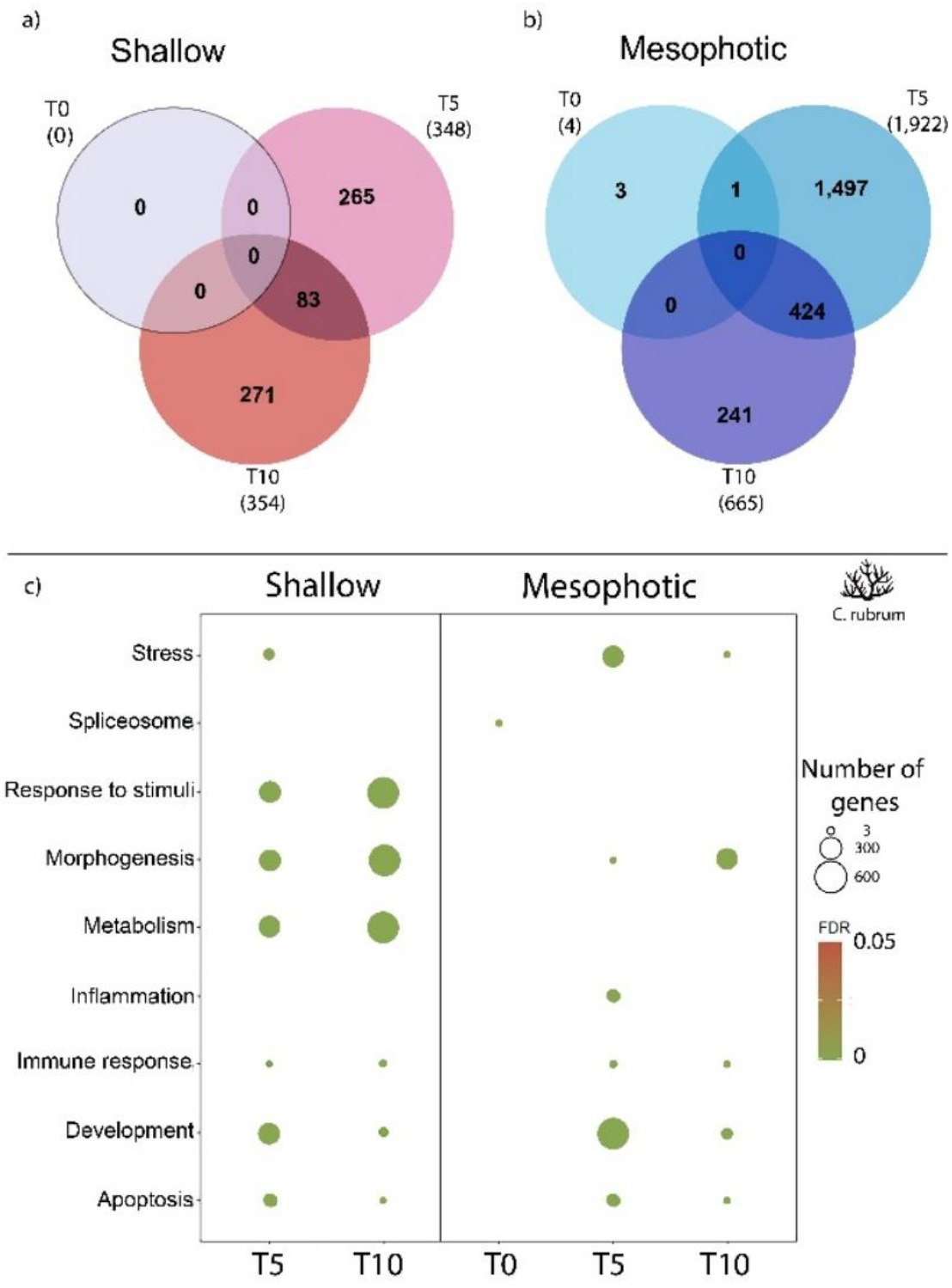
Number of differentially expressed genes (DEGs) between control and stress by experimental days (T0, T5, and T10) and their common overlap in a) Shallow and b) Mesophotic populations of *C. rubrum*. c) Significant functional enrichment of Gene Ontology (GO) terms across the entire experiment for each population. No enrichment was found for T0 at the Shallow population. The size and the color of the circles are proportional to the number of genes observed within each GO category and to their significance, respectively (FDR value< 0.05)

For the mesophotic population, we observed that at T0, DEGs related to transcription, RNA binding and metabolism were underlying three enriched functions concerning the spliceosomal complex processes. (*At5g10370, Isy1*, Table S7a, b). Moreover, at T5, 1,497 DEGs altered their transcription and represented the largest changes in gene expression across days for this population (863 upregulated, 634 downregulated; Figure 4b, c, Table S7a). Processes of development and stress significantly characterize thermal responses at this time point with DEGs related to growth factor receptors (*FGFR3*) bone remodeling (*CTSK*), collagen metabolism (*ADIPOQ*), ROS species (*BBOX1*), heat shock proteins (*HSPA5*), cellular response to environmental stress (*Mapk8, GRB7*) and inflammation (*CTSL*) (Table 7a, c). Finally, a variety of cellular structure processes were also enriched underlined by histones (*ABCB4*) and microtubules genes (*MYOF*) (Table S7c). At last, at T10, the mesophotic population differentially expressed 241 unique genes (60 upregulated, 181 downregulated) with the majority of them related to morphogenesis processes mediated by collagen (*MMP14*), elastin (*Mmp2*), transcription factors (*Uncx*) and innate immunity (*Rnasel*) (Figure 4c, Table S7d). In particular, DEGs related to heat shock proteins chaperones (*HSPA2*), Aquaporins (*AQP3*), inflammation (*PARP14, TRPA1*), transcription factor (*IRF2*), collagen (*Mmp2*) and several energy processes (*PRKAG2, glul, heix*) characterized the stress signatures during this day. Only two GO terms were related to stress (GO:0033554: cellular response to stress) and apoptosis (GO:0043065: positive regulation of apoptotic process) with further DEGs related to embryogenesis (*AFDN, RERG*), ion transport (*Slc4a3, cho-1*), myosin (*MYL9, Mmp2*) and metabolism (*Megf10, MMP14, LARP4B*) (Table S7d). While, GO terms related to immune responses were particularly underlined by transcription factors (*IRF2*), immune response genes (*Clec4g, CTSC*), heat shock proteins (*HSPA2*), and inflammation (*PARP14*). Interestingly, *CTSC* and *HSPA2* were downregulated and presented the largest changes in expression (log2foldchange<-20) in comparison with control conditions (Table S7a, d).

### Frontloaded gene expression

We compared the gene expression levels in control conditions of shallow vs mesophotic populations for those genes that were differentially expressed in the mesophotic population (i.e. upregulated genes, Table S8a). Thus, of the total 882 unique DEGs from the mesophotic population, 310 DEGs were found upregulated during thermal stress (Table S8a). Among these 310 DEGs, 172 were found frontloaded, meaning that they were already upregulated in the shallow population in the control treatment. In addition, of the total set of 310 DEGs, 120 were classified as “greater fold change” and 6 as “reduced reaction” (Table S8a). The 12 DEGs left were not significantly differentially expressed either in the shallow population at control or at stress conditions, making it difficult to assess their status. As such, they were correspondingly classified as “NA” (Table S8a). From the 172 frontloaded DEGs, 134 had a valid annotation and suggested frontloaded cellular responses, apoptosis, immune responses, response to hypoxia and starvation, as well as functions related to RNA transcription (Table S8b).

## DISCUSSION

In this study, we revealed significant differences in the early transcriptomic responses to thermal stress in two populations of *Corallium rubrum* coming from a shallow and a mesophotic environment. Overall, the thermal stress (>25°C) generates significant alterations of the transcriptomic responses, especially at day 5, in the two populations. However, the shallow population exhibits a more subdued response compared to the mesophotic population, with fewer DEGs, distinct molecular functions involved, a distinct temporal pattern of gene expression and a frontloaded expression of various genes (e.g. apoptosis). We discuss how these patterns are likely driven by local adaptation to thermal conditions and their implications for the species’ conservation.

### Conserved heat shock proteins and regulatory pathways drive early stress responses in C*orallium rubrum*

Based on the entire dataset (control samples vs stress samples), we first demonstrate how the early responses to thermal stress in *C rubrum* rely on the expression of heat shock proteins (HSPs) and transcription factors, suggesting diversified responses of cellular stress. The expression of HSPs likely contributes to maintaining overall coral health and promoting heat acclimation, as they represent a primary protective transcriptomic response to stress in tropical hexacorals like *Acropora* spp. (Desalvo et al. 2008; Barshis et al. 2013; Palumbi et al. 2014; Seneca and Palumbi 2015; Traylor-Knowles et al. 2017b) and reflect an evolutionarily conserved mechanism among cnidarians (Black et al. 1995; Hayes and King 1995; Downs et al. 2000; Brown et al. 2002; Robbart et al. 2004, 2004b; Rossi et al. 2006; Rosic et al. 2011). Concomitantly, the expression of transcription factors like *Atf6*, and *AP-1*, known to be associated with bleaching responses in tropical hexacorals (Bellantuono et al. 2012; Ruiz-Jones and Palumbi 2017; Traylor-Knowles et al. 2017b, 2017a), also suggests that such genes may regulate early cellular activity related to tissue integrity in *C. rubrum*, since it is an aposymbiotic octocoral that develops tissue necrosis in response to MHWs (Garrabou et al. 2001). Besides HSPs and transcription factors, the thermal stress induced further gene expression consistent with the “core cnidarian heat stress response” (Cziesielski et al. 2018) associated with regulation of heat stress genes (*HSPA6, HSP90B1, AP1M1, Atf6*), and major antioxidant genes (i.e. ROS species, *SOD, YqjG*), calcium (Ca^2+^) homeostasis (*Calmodulin, Cib2*), extracellular matrix restructuration (*MMP14, Adam10, SRPRA*), cytoskeleton rearrangements (*FCN2, ACT1, Arhgef17*), apoptosis (*TRAF2, 4, Mapk14*) and ribosomal genes (*RPLP0, 4, 13a, 18, mrpl52*). This expression of the “core cnidarian heat stress response” point towards an evolutionarily conserved response to thermal stress shared among temperate octocorals and tropical hexacorals.

### Differential expression patterns and wound healing responses suggest increased thermal sensitivity in mesophotic populations

When considering control and treatment samples independently for each population, we highlighted variability in early transcriptomic responses to thermal stress between shallow and mesophotic populations. Overall, the shallow population displayed significantly less transcriptional modifications with lower thermal stress signatures (*i*.*e*. number of DEGs and GO-terms) compared to the mesophotic population. This was in concordance with previous studies in *C. rubrum* suggesting a larger number of transcripts (i.e. overexpression) for populations from 40m (120 contigs) vs 5m (92 contigs) (Pratlong et al. 2015). Moreover, in the mesophotic population, GO-terms related with cell division, reactive oxygen species (ROS), cell death, protein degradation, NF-Kappa B signaling, immune responses and protein folding were identified. These broader biological alterations are in concordance with the generalized coral environmental stress response (ESR) proposed by Dixon et al., 2020, and more specifically with the ESR-type A. The ESR-type A is characterized by the upregulation of transcriptional processes of cell death, ROS species, NF-kappa B signaling, immune responses and protein folding genes, indicating a more severe stress response compared to the ESR-Type B which implies a downregulation of these processes and to a less severe stress response. Thus, on one hand, our results highlight the same upregulated transcriptional processes characteristic of this type of stress response (i.e., ESR-Type A). At the same time, the greater number of differentially expressed transcripts in the mesophotic population suggests broader stress severity and heightened sensitivity, consistent with observations in other cnidarians (i.e. *Acropora* spp; Dixon et al., 2020). In addition, such differential responses to thermal stress expanded and complemented previous studies conducted in the red coral (Torrents et al. 2008; Haguenauer et al. 2013; Ledoux et al. 2015; Pratlong et al. 2015), and in other coral species (Crisci et al. 2017; Franzellitti et al. 2018; Studivan and Voss 2020). Particularly, *C. rubrum* populations from mesophotic habitats have shown a pronounced vulnerability to thermal stress with higher levels of tissue necrosis (phenotypic level) (Torrents et al. 2008; Ledoux et al. 2015) and contrasted patterns of gene expression (molecular phenotypic level) compared to shallow populations, highlighting the expression of *HSP70* as one of the main adaptations to shallow habitats (Haguenauer et al. 2013). Although we did not observe *HSP70* transcription, the shallow samples upregulated *HSP90B1* during thermal stress treatment, which acts as a chaperone generating ATP-dependent cellular processes during heat shock treatments (Zhang et al. 2020), similar to *HSP70*. Likewise, the upregulated expression of *ST13* (cytosolic Hsp70-interacting protein Hip) which is known for interacting with *HSP70* to stabilize protein complexes and maintaining homeostasis (Li et al. 2013), was observed on day 10^th^ in the shallow population, suggesting protein folding repair regulation at the end of the experiment. Regarding the mesophotic population, its sensitivity to thermal stress was further suggested by the unique expression of “wound healing” mechanisms, which are characterized mainly by the expression of collagen and previously reported in reaction to heat stress (Desalvo et al. 2008; Barshis et al. 2013; Traylor-Knowles 2019). We found several up and downregulated DEGs and GO terms related to collagen, myosin and fibrillin in this population which may maintain structural integrity of the extracellular matrix and an initiation of cascades of healing factors due to stress (Traylor-Knowles 2019). Mechanisms of tissue resize and recovery have been observed in the red coral (Roveta et al. 2023) as well as in octocoral *Heliopora coerulea* (Orgel et al. 2017; Guzman et al. 2019) and in *A. hyacynthus* (Traylor-Knowles 2019). Accordingly, the alteration of wound healing mechanisms observed may serve as an immediate molecular response of protection from heat stress in the mesophotic population.

### Distinct temporal trajectories of thermal stress response in *C. rubrum*

We then compared the dynamics of gene expression across experimental days between the two populations. Stronger and contrasting transcriptional transitions were observed at days 5^th^ and 10^th^ for the mesophotic and shallow population. In particular, a large transcription of “developmental” processes (i.e. muscle structure development, tissue morphogenesis) and DEGs associated with bone development (*SOX9, Lrp6*), ubiquitin tumor repression (*mdm2, FHL3, NEO1*), tissue and organ formation (*TWIST2, SIX6, Uncx, Notch1*) suggested that the activation of tissue alterations signaling cascades (Traylor-Knowles et al. 2017b) was critical in the last part of the experiment for the shallow population (day 10^th^). In contrast, larger transcriptional changes were evident earlier at day 5^th^ for the mesophotic population. In this latter case, a different set of DEGs and GO-terms functions (i.e. organ development) associated with lipids (*Git2, ABCB4, Musk, Map3k9*), recycling of immune signaling components (*ift20*) and inflammation (*Btrc, CATL*), suggested that transcription of immune and inflammation pathways were induced early in this population to respond to heat stress and maintained individual integrity (Anderson et al. 2016). Accordingly, we revealed not only a distinctive timing in the activation of thermal stress mechanisms (early in mesophotic individuals - day 5^th^, and late in shallower individuals - day 10^th^), but also a specific set of molecular functions and potential adaptation mechanisms in each population to counteract the same heat stress.

### First evidence of frontloading in thermal stress response

Our results support, for the first time, the occurrence of transcriptional frontloading in the red coral. Transcriptional frontloading received particular attention in recent years since it enables corals to preemptively activate stress response pathways and enhance their resilience to environmental stressors (Barshis et al., 2013; Brener-Rafalli et al., 2022; Collins et al., 2021). Here, the frontloading strategy relies on the expression of HSPs (*HSPA5* and *9*), redox-active proteins (*TXN, CYP169*), apoptosis regulation (*Casp9, PARP12, 14, PIAP, Litaf*) and collagen genes (*Loxl2, COL4A3*) which suggested potential molecular predispositions of the shallow population to respond to heat stress, including, for instance, shifts of cellular set points at which apoptosis activation occur (Ainsworth et al. 2011; Barshis et al. 2013; Pratlong et al. 2015; Maor-Landaw and Levy 2016). Accordingly, these frontloaded genes are likely key processes allowing the shallow population to better withstand with heat stress without extensive transcriptomic reprogramming as observed in the mesophotic population. Further studies are needed to test whether or not frontloading gene expression is ubiquitous among shallow red coral populations.

### Local adaptation as a potential driver of thermal stress responses in *C. rubrum*

Based on the qualitative and quantitative differences, the contrasting temporal dynamics of gene expression between the two populations, and the frontloaded gene expression observed in the shallow population, we hypothesize that each population has potentially evolved local adaptations to their respective environments, influencing their capacity to withstand and respond to heat stress. Local adaptation to thermal environment has been suggested previously as a main process underlying the differential responses to thermal stress among shallow and mesophotic populations in *C. rubrum* using *in situ* common garden experiment (Ledoux et al. 2015) and genomic approaches (Pratlong et al. 2021). The shallow and mesophotic populations are separated by a 4 km distance, a spatial distance at which significant genetic structure was demonstrated (Horaud et al. 2024). In addition, the thermal regimes between shallow and mesophotic habitats differed markedly (Bensoussan et al. 2019, Fig 1). The shallow population is exposed to greater temperature fluctuations and experiences higher peak maximum summer temperatures compared to the mesophotic population. Accordingly, we posit that the fine-scale genetic divergence combined with contrasted local environmental conditions has potentially driven local adaptation which may in turn influence gene expression profiles in response to thermal stress. Indeed, gene expression regulation is highly heritable and evolves in response to selective pressures between environments (Whitehead and Crawford 2006). Thus, the delayed yet robust transcriptional shifts in the shallow population may reflect an adaptive strategy to cope with the higher temperature conditions driving larger tolerance to thermal stress. Similar local adaptation patterns have been elucidated into other coral species and several other taxa of plants and invertebrates (Hereford 2009; Kenkel and Matz 2016). For instance, shallow populations in different *Acropora* species show specific gene expression patterns, which enhance resilience to the extreme conditions (Barshis et al. 2013; Ruiz-Diaz et al. 2022; Thomas et al. 2022).

### Conservation implications for red coral populations

The profound differences in the early transcriptomic responses to thermal stress between shallow and mesophotic populations of *C. rubrum* underscore that these populations rely on fundamentally distinct molecular strategies in both environments. This has important conservation implications. The deep refugia hypothesis proposes that mesophotic populations serve as a reservoir of genetic diversity and potential sources of recruits for shallow populations (Glynn 1996). This has been observed after severe El Niño events (1982–1983 and 1997–1998) with rapid recolonization of depleted shallow populations of *Millepora intricata* (<12m) from surviving deep-water colonies (>11m; Smith et al. 2014). In the present case, the distinct transcriptomic signatures observed in the mesophotic population under thermal stress suggest that these individuals may have limited adaptive potential to persist in shallower habitats increasingly affected by marine heatwaves. Exposure to naturally warmer and more extreme thermal regimes in shallow waters may be detrimental to recruits originating from mesophotic populations and also raises concerns about the effectiveness of using mesophotic populations as sources for active restoration efforts in these shallow environments. While these concerns align with previous studies (e.g., Ledoux et al., 2015), recent evidence has demonstrated the long-term (i.e., over 10 years) effectiveness of active transplantation of *C. rubrum* colonies (Zentner et al., 2025). Further studies are needed to reconcile these findings and better understand the conditions under which transplantation can contribute to shallow-water active restoration efforts.

## Conclusion

In conclusion, our findings offer insights into potential evolutionary processes that enable populations to cope with ongoing changing environments that should inform conservation and restoration strategies. Further research should explore the heritability of thermal responses to better assess the adaptive potential and resilience of *C. rubrum* under climate change. Additionally, analyzing life history traits such as survival, growth, and reproduction will be crucial to elucidate potential fitness trade-offs inherent to local adaptation to thermal environment. This will provide a more complete picture of the evolutionary constraints and adaptive potentials of Mediterranean octocorals, ultimately guiding timely and more effective conservation and restoration strategies.

## Data Availability Statement

The data and code that support the findings of this study are openly available in the GitHub repository https://github.com/sandrarcr/Corallium_rubrum_RNAseq_local_adaptation (raw counts data, scripts for DESeq2 analysis and associated plots) and in Zenodo data repository at https://doi.org/10.5281/zenodo.15388872. Temperature data records are publicly available in Zenodo at https://doi.org/10.5281/zenodo.14007188 and in GitHub at https://github.com/Damyck/tMednet.git (SST data). Raw reads for RNAseq information are in process for release in the European Nucleotide Archive (ENA).

## Statements and Declarations Acknowledgments

The authors would like to thank the technician staff at the Institute of Marine Sciences, Barcelona, for their support with the experimental setup, to the members of MedRecover research group from the “Generalitat de Catalunya” for supporting fieldwork campaigns and to the CIIMAR Biodiscovery & Omics technology platform. CNAG acknowledges the support of the Spanish Ministry of Science and Innovation through the Instituto de Salud Carlos III and the 2014–2020 Smart Growth Operating Program and co-financing with the European Regional Development Fund (MINECO/FEDER, BIO2015-71792-P). We also acknowledge the support of the Generalitat de Catalunya through the Departament de Salut and Departament d’Empresa i Coneixement.

## Funding

This work was financially supported by the European Union’s Horizon 2020 research and innovation programme (SEP-210597628—FutureMARES), the Interreg EuroMed MPA4Change (EURO-MED0200736) and the project CORFUN (TED2021-131622B-I00). This research has also been funded by the Short-Term Scientific Mission (STSM) grant provided by the COST Action CA20102-MAF World (E-COST-GRANT-CA20102-f04dde3c & E-COST-GRANT-CA20102-a33d8dce). SRC was supported by the scholarship Doctorados en el exterior No. 906 de 2021 of Ministerio de Ciencia, Tecnología e Innovación of Colombia. JBL was supported by the strategic funding UIDB/04423/2020, UIDP/04423/2020 and 2021.00855.CEECIND through national funds provided by FCT -Fundaço para a Ciência e a Tecnologia. SRC, JG, MJ, MZ, PLS and JBL are part of the Marine Conservation research group - MedRecover (2021 SGR 01073) from the “Generalitat de Catalunya”. SRC, JG, MJ and PLS acknowledge grant CEX2024-001494-S funded by AEI 10.13039/501100011033.

## Author information

### Contributions

**Sandra Ramirez-Calero**: conceptualization, data curation, data analysis, funding acquisition, writing – review and editing original draft. **Joaquim Garrabou**: conceptualization, data collection, experimental setup, data analysis, review and editing, funding acquisition. **Sneha Suresh**: data analysis validation, review and editing. **Marta Gut**: sequencing, review and editing. **Marc Jou**: data visualization, review and editing. **Xenia Sarropoulou**: laboratory analysis and extractions, review and editing. **Paula Lopez-Sendino**: data collection, experimental setup, laboratory analysis and extractions, review and editing. **Mikel Zabala**: data collection, review and editing. **Jean-Baptiste Ledoux**: conceptualization, data collection, experimental setup, laboratory analysis, data analysis, write – review and editing, funding acquisition.

## Conflict of interest

The authors have no relevant financial or non-financial interests to disclose.

## Ethics declarations

All applicable international, national, and/or institutional guidelines for animal testing, animal care, and use of animals were followed by the authors.

## Notes

https://github.com/sandrarcr/Corallium_rubrum_RNAseq_local_adaptation

https://github.com/marcjou/tMednet

### Competing Interest Statement

The authors have declared no competing interest.

