## Supplementary Material 1 for "Does local adaptation influence thermal responses in red coral populations across depth gradients? Transcriptomic insights for effective conservation"

Journal name: Coral reefs

Ramirez-Calero, S.[1,2*], Garrabou J.[1] , Suresh S.[3], Gut, M[4]., Jou, M[1]., Sarropoulou, X.[5], Lopez-Sendino, P.[1], Zabala, M.[2], Ledoux, JB[5+].

[1] Institute of Marine Sciences, ICM-CSIC, Pg. Marítim de la Barceloneta 37-49, 08003, Barcelona, Spain

[2] Facultat de Biologia, Universitat de Barcelona, Av. Diagonal 643, 08028, Barcelona, Spain.

[3] Department of Environmental Conservation, University of Massachusetts, Amherst, MA 01003, USA

[4] Centro Nacional de Análisis Genómico (CNAG), Barcelona 08028, Spain

[5] CIIMAR/CIMAR LA, Centro Interdisciplinar de Investigação Marinha e Ambiental, Universidade do Porto, Terminal de Cruzeiros do Porto de Leixões, 4450-208 Matosinhos, Portugal.

In this Supplementary Information, we provide further details regarding:

- *Sampling and experimental setup:* Aquarium setup and common garden experiment conditions (Figure S1)
- *Thermal regime characterization for Shallow and Mesophotic sites:* Mean temperature values and Standard Deviation (SD) 2002-2024 (Figures S2a, b)
- De novo *transcriptome assembly:* Transcriptome quality assessement and BUSCO results (Figure S3)
- *Differential gene expression analyses:* Principal component analysis (PCA, Figure S4)

*Sampling and experimental setup*

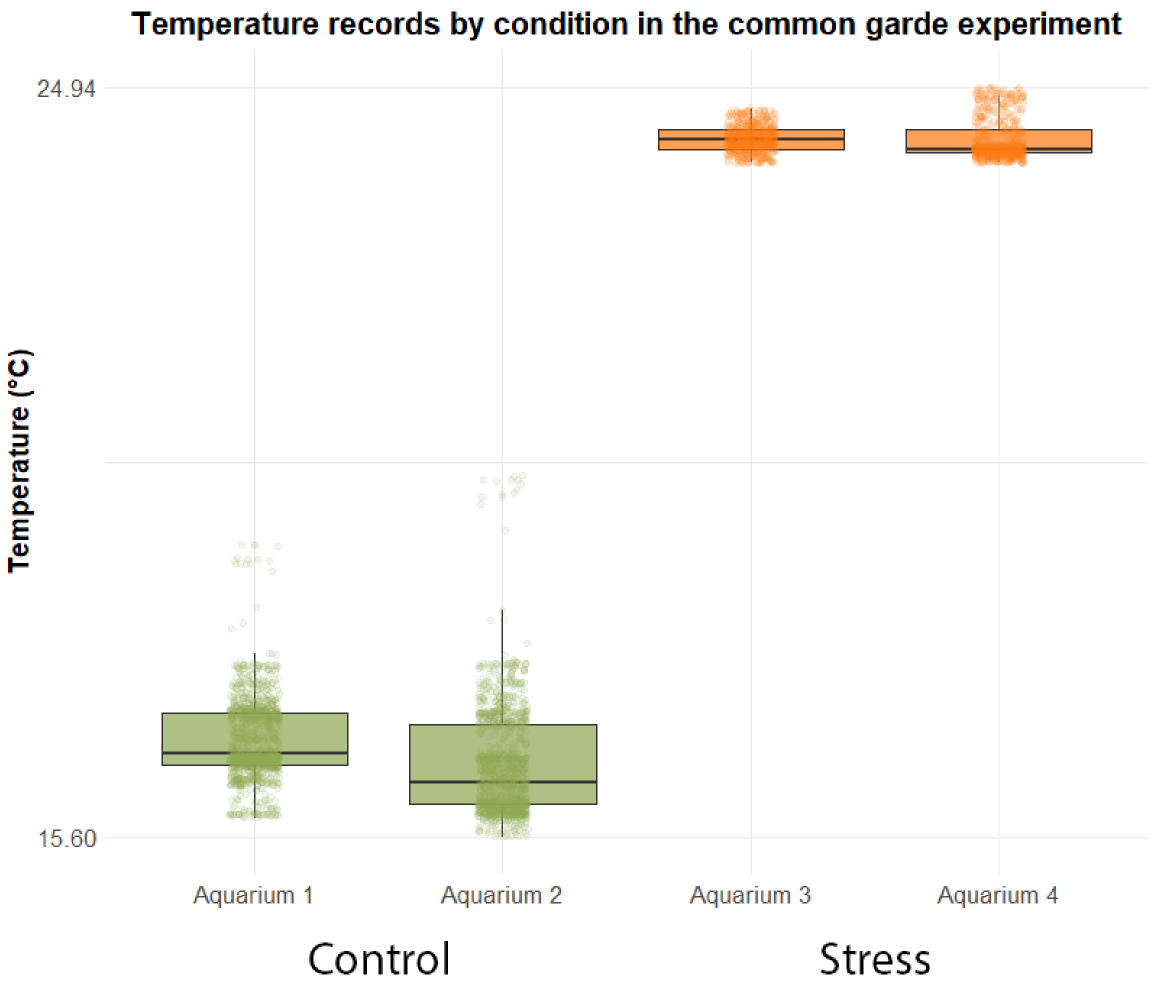


**Figure S1.** Boxplot representing seawater temperature in aquariums for the common-garden experimental setup for individuals of shallow and mesophotic populations of *Corallium rubrum* exposed to two treatments: i) control (green-18°C), and ii) thermal stress (orange-25°C,).

*Thermal Regimes characterization for Shallow and Deep sites*

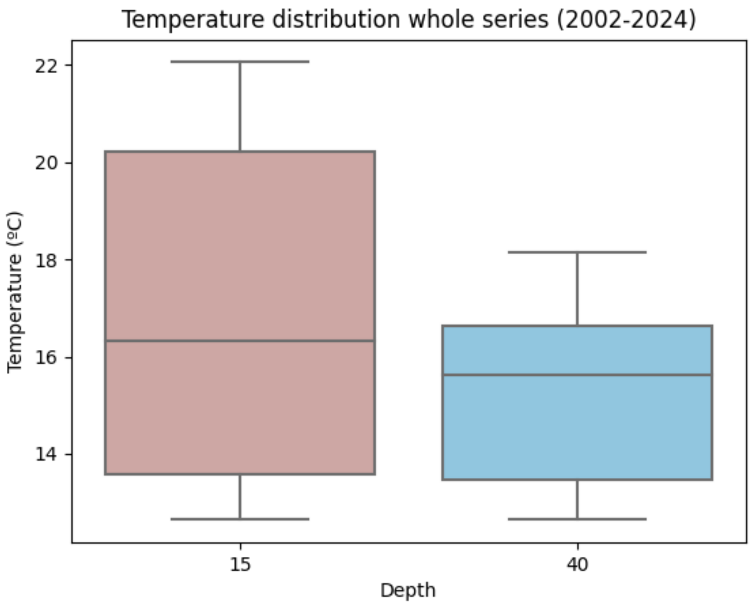


**Figure S2a**. Mean temperatures boxplots from the entire series record (2002-2024) in Shallow (pink-15m) and Mesophotic (blue-40m) populations using the T-MEDNet database (Benssousan et al. 2019).

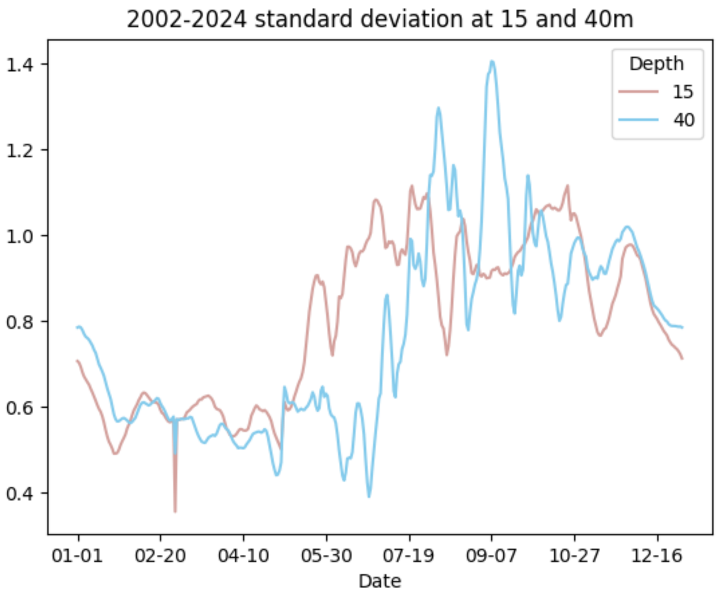


**Figure S2b.** Standard Deviation (SD) for mean temperature values in Shallow (pink-15m) and Mesophotic (blue-40m) sites from the entire series record (2002-2024). SD values are based on the filtered data set to reduce inconsistencies. The resultant gaps in some temperature records created some of the spikes that can be seen in the graph. Thus, the filtering was treated within blocks so distant dates remain disconnected.

*De novo Transcriptome assembly*

The total number of Trinity genes was 293,131 while transcripts were 533,146 in total. Both represent the full set of assembled transcripts, while the GC content of 40.05% reflects the base composition of the assembled sequences (Table S2a-c). In addition, Assembly statistics based on all transcript contigs revealed that the median contig length was 357 bp, and the average contig length is 624.22 bp. Assembly statistics based only on the longest isoform per gene revealed similar values to all transcript contigs with a median contig length was 316 bp, and the average contig length was 525.95 bp (Table S2b, c). The N10 through N50 values were computed based on all assembled contigs. As such, 10% of the assembled bases were found in transcript of at least 3,669 bp in length (N10 value), and the N50 value indicated that at least half of the assembled bases were found in contigs of at least 895 bases in length.

Bowtie statistics revealed that the quality of the alignment and the effectiveness of the assembly for each sample were generally consistent, with overall alignment rates ranging from 67% to 72%. The alignment rates varied slightly between the samples, with most samples showing an alignment rate close to 70% (Table S2d).

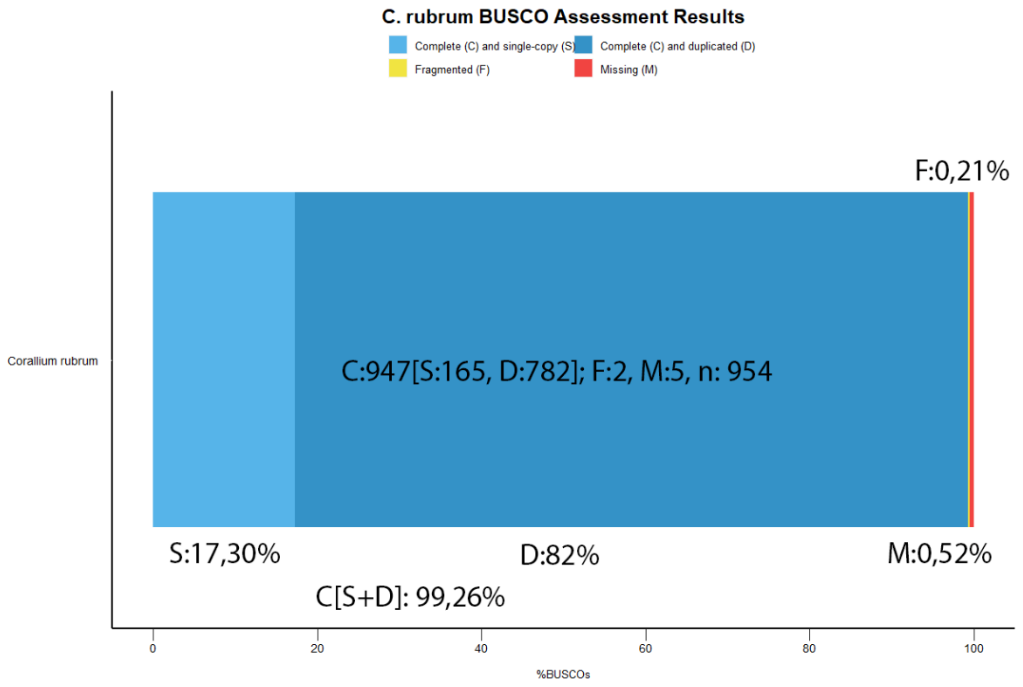


**Figure S3.** BUSCO plot showing the completeness of the *de novo* transcriptome assembly of *Corallium rubrum* based on conserved orthologous gene sets using *metazoan_odb10* database. Categories are found at the top of the graph.

*Differential gene expression analyses*

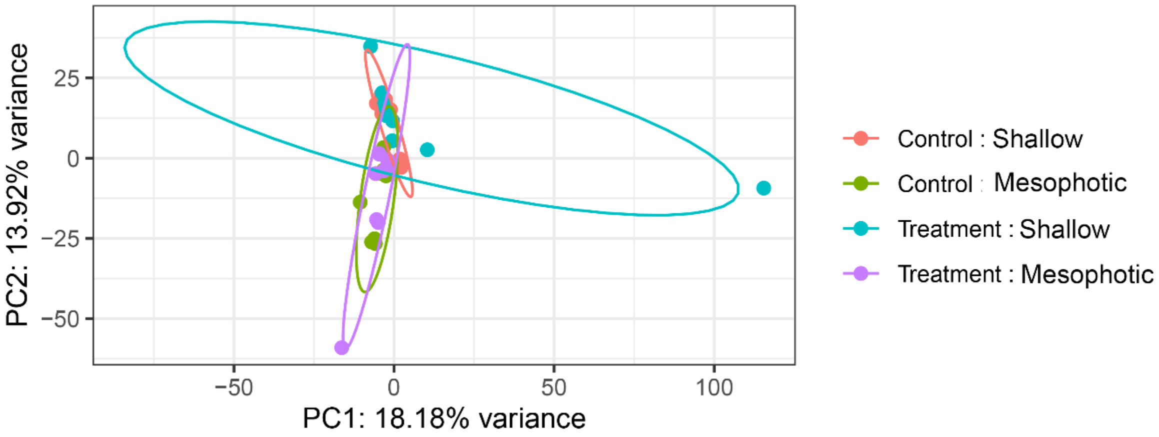


**Figure S4.** Principal Component Analysis (PCA) of overall transcript expression profiles across all samples. The analysis was conducted to assess sample clustering and detect potential outliers between two environmental conditions: Control and Thermal stress. Each point represents a sample, colored by group. PC1 explains 18.18% of the variance and PC2 explains 13.92%. This plot confirms the consistency within each condition and highlights overall transcriptomic differences between shallow and deep samples.
